# Structural basis for the inhibition of SARS-CoV-2 main protease by antineoplastic drug Carmofur

**DOI:** 10.1101/2020.04.09.033233

**Authors:** Zhenming Jin, Yao Zhao, Yuan Sun, Bing Zhang, Haofeng Wang, Yan Wu, Yan Zhu, Chen Zhu, Tianyu Hu, Xiaoyu Du, Yinkai Duan, Jing Yu, Xiaobao Yang, Xiuna Yang, Kailin Yang, Xiang Liu, Luke W. Guddat, Gengfu Xiao, Leike Zhang, Haitao Yang, Zihe Rao

## Abstract

The antineoplastic drug Carmofur was shown to inhibit SARS-CoV-2 main protease (M^pro^). Here the X-ray crystal structure of M^pro^ in complex with Carmofur reveals that the carbonyl reactive group of Carmofur is covalently bound to catalytic Cys145, whereas its fatty acid tail occupies the hydrophobic S2 subsite. Carmofur inhibits viral replication in cells (EC_50_ = 24.30 μM) and it is a promising lead compound to develop new antiviral treatment for COVID-19.

---

COVID-19, a highly infectious viral disease, has spread since its appearance in December 2019, causing an unprecedented pandemic. The number of confirmed cases worldwide continues to grow at a rapid rate, but there are no specific drugs or vaccines available to control symptoms or the spread of this disease at this time.

The etiological agent of the disease is the coronavirus SARS-CoV-2. This virus has a ∼30,000 nt RNA genome. The N-terminus of the viral genome encodes two translational products, polyproteins 1a and 1ab (pp1a and pp1ab)^1,2^, which are processed into mature non-structural proteins, by the main protease (M^pro^) and a papain-like protease^3^. M^pro^ has been proposed as a therapeutic target for anti-coronavirus (CoV) drug development^4-6^. We previously screened over 10,000 compounds and identified Carmofur as compound that can inhibit M^pro^ *in vitro*, with an IC_50_ of 1.82 μM^7^.

Carmofur (1-hexylcarbamoyl-5-fluorouracil) is a derivative of 5-fluorouracil (5-FU) (Fig. 1a) and an approved antineoplastic agent. Carmofur has been used to treat colorectal cancer since 1980s^8^, and has shown clinical benefits on breast, gastric, and bladder cancers^9-11^. The target for Carmofur is believed to be thymidylate synthase^12,13^, but it has also been shown to inhibit human acid ceramidase (AC)^14^, through covalent modification of its catalytic cysteine^15^.

**Fig. 1.**
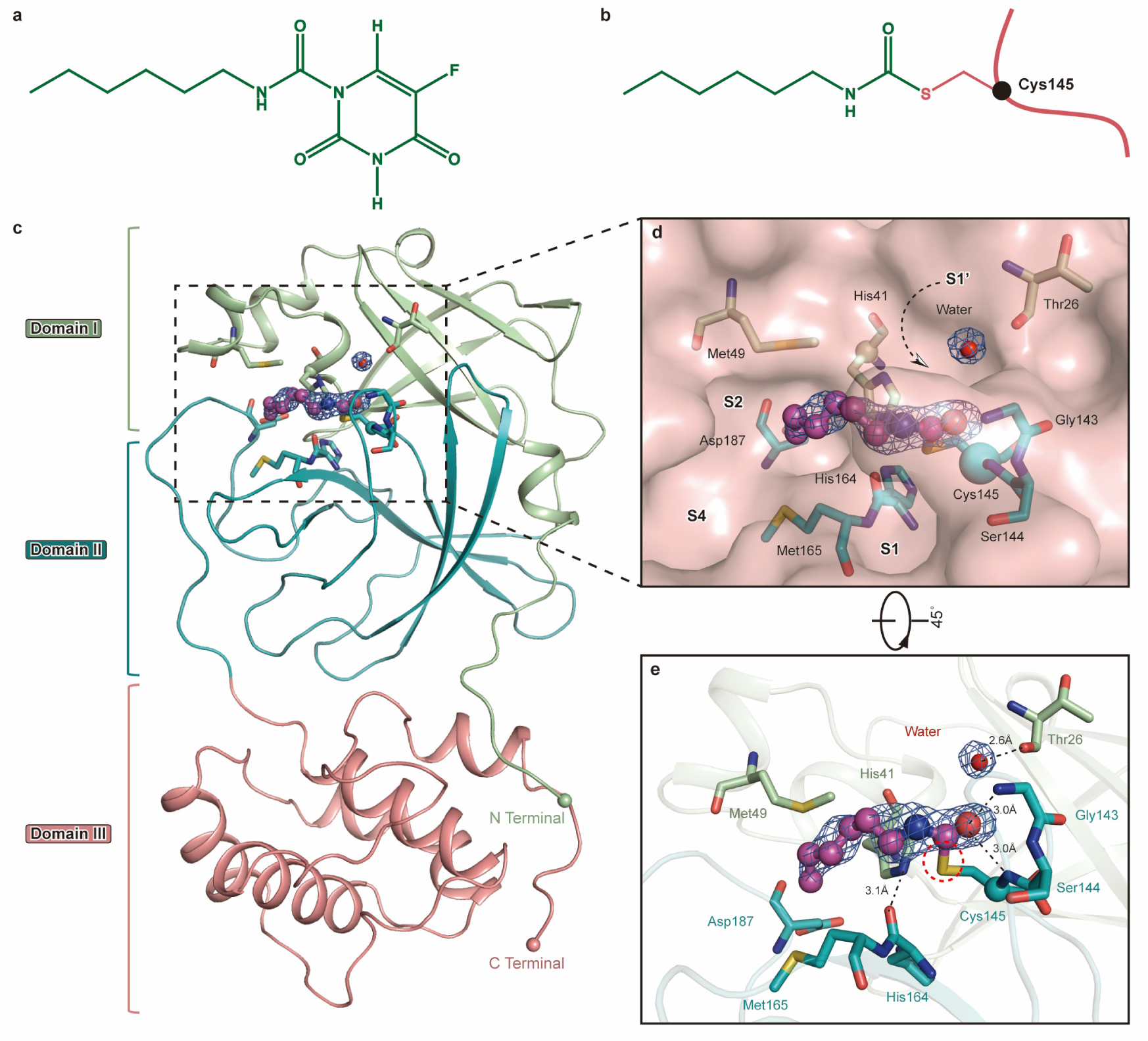
SARS-CoV-2 M^pro^ in complex with Carmofur. **a**, The chemical structure of Carmofur. **b**, The binding mode of Carmofur to SARS-CoV-2 M^pro^. The red curve represents SARS-CoV-2 M^pro^ polypeptide with the sidechain of Cys145 protruding. **c**, The structure of a single protomer. The three domains are shown in three different colors. The catalytic center is located within the dashed square. **d**, Zoom in of the catalytic center. The residues that participate in Carmofur binding are shown as stick models. Carmofur is show as a ball-and-stick model with the carbons in magenta. Water is presented as a red sphere. **e**, A rotated view of the binding site, but with the surface removed. The red dashed circle highlights the C-S covalent bond.

The molecular details for how Carmofur inhibits M^pro^ activity were unresolved. Here, we present the 1.6 Å X-ray crystal structure of SARS-CoV-2 M^pro^ in complex with Carmofur (Fig. 1b, c and Supplementary Table 1). In agreement with previous studies^4,5,16-18^, M^pro^ forms a homodimer (protomer A and B) related by crystallographic symmetry (Extended Data Fig. 1a, b). All of the residues (1–306) in the polypeptide could be traced in the electron density map. Each protomer is composed of three domains (Fig. 1c): domain I (residues 10–99), domain II (residues 100–184), and domain III (residues 201–303); and a long loop region (residues 185–200) connects domains II and III. The substrate-binding pocket lies in the cleft between domain I and domain II feature the catalytic dyad residues Cys145 and His41 (Fig. 1c, d). The substrate-binding pocket is divided into a series of subsites (including S1, S2, S4, and S1′), each accommodating a single but consecutive amino acid residue in the substrate. The first residue serine of one protomer interacts with residue Phe140 and Glu166 of the other protomer to stabilize the S1 subsite (Extended Data Fig. 1c), and this structural feature is essential for catalysis^7^.

The electron density map unambiguously shows that the fatty acid moiety (C_7_H_14_NO) of Carmofur is linked to the Sγ atom of Cys145 through a 1.8 Å covalent bond, whereas the fatty acid tail is inserted into the S2 subsite (Fig. 1d, e). This observation suggests that the sulfhydryl group of Cys145 attacks the electrophilic carbonyl group of Carmofur, resulting in covalent modification of the cysteine residue and release of the 5-fluorouracil moiety (Fig. 1b and Extended Data Fig. 2a). In addition to the C-S covalent bond (highlighted in Fig. 1e), the inhibitor is stabilized by numerous hydrogen bonds and hydrophobic interactions (Fig. 1e and Extended Data Fig. 2b). The carbonyl oxygen of Carmofur occupies the oxyanion hole and forms hydrogen bonds (3.0 Å) with the backbone amides of Gly143 and Cys145, mimicking the tetrahedral oxyanion intermediate formed during protease cleavage (Fig. 1e). The fatty acid tail, which appears in an extended conformation, inserts into the bulky hydrophobic S2 subsite (composed of the side chains of His41, Met49, Tyr54, Met165, and the alkyl portion of the side chain of Asp187) (Fig. 1d, e). The hydrophobic interactions are mainly contributed by the side chains of His41, Met49 and Met165, all of which run parallel with the alkyl part of the fatty acid tail of the inhibitor (Fig. 1e and Extended Data Fig. 2b).

The mechanism of covalent modification by Carmofur is different from that of the inhibitor N3^7,19^, which covalently modifies Cys145 through Michael addition of the vinyl group. The structures of M^pro^-Carmofur and M^pro^-N3 are overall similar (r.m.s.d. of 0.286 Å for all C*α* atoms). The largest conformational differences occur in the substrate binding pocket, with the backbone surrounding Carmofur in a slightly more outward position compared with the M^pro^-N3 complex structure (Extended Data Fig. 3a). Another difference is that Carmofur only occupies the S2 subsite (Fig. 1d), whereas N3 occupies four subsites (S1, S2, S4 and S1′, see Extended Data Fig. 3b, c). The lactam ring of N3 is located in the S1 subsite, which is filled by a DMSO molecule in the M^pro^-Carmofur structure (Extended Data Fig. 3b, c). These observations demonstrate the potential for structural elaboration of Carmofur and will be useful to design more potent derivatives against the M^pro^ of SARS-CoV-2.

We previously showed that treatment with 10 μM Ebselen (EC_50_ = 4.67 μM) inhibited infection of Vero cells with SARS-CoV-2 whereas Carmofur did not showed detectable antiviral activity at this concentration^7^. Here we determined the inhibitory effect of Carmofur against SARS-CoV-2 infection on Vero E6 cells, as previously described^20^ (Fig. 2). By measuring viral RNA in supernatant, we determined the EC_50_ for Carmofur as 24.30 μM (Fig. 2a). To verify this result, we fixed infected cells and stained them using anti-sera against viral nucleocapsid protein (NP) and observed a decrease in NP levels after Carmofur treatment (Fig. 2b). We also performed cytotoxicity assays for Carmofur in Vero E6 cells and determined the CC_50_ value of 133.4 μM (Fig. 2c). Thus, Carmofur has a favorable selectivity index (SI) of 5.36, but further optimization will be required to develop into an effective drug.

**Fig. 2.**
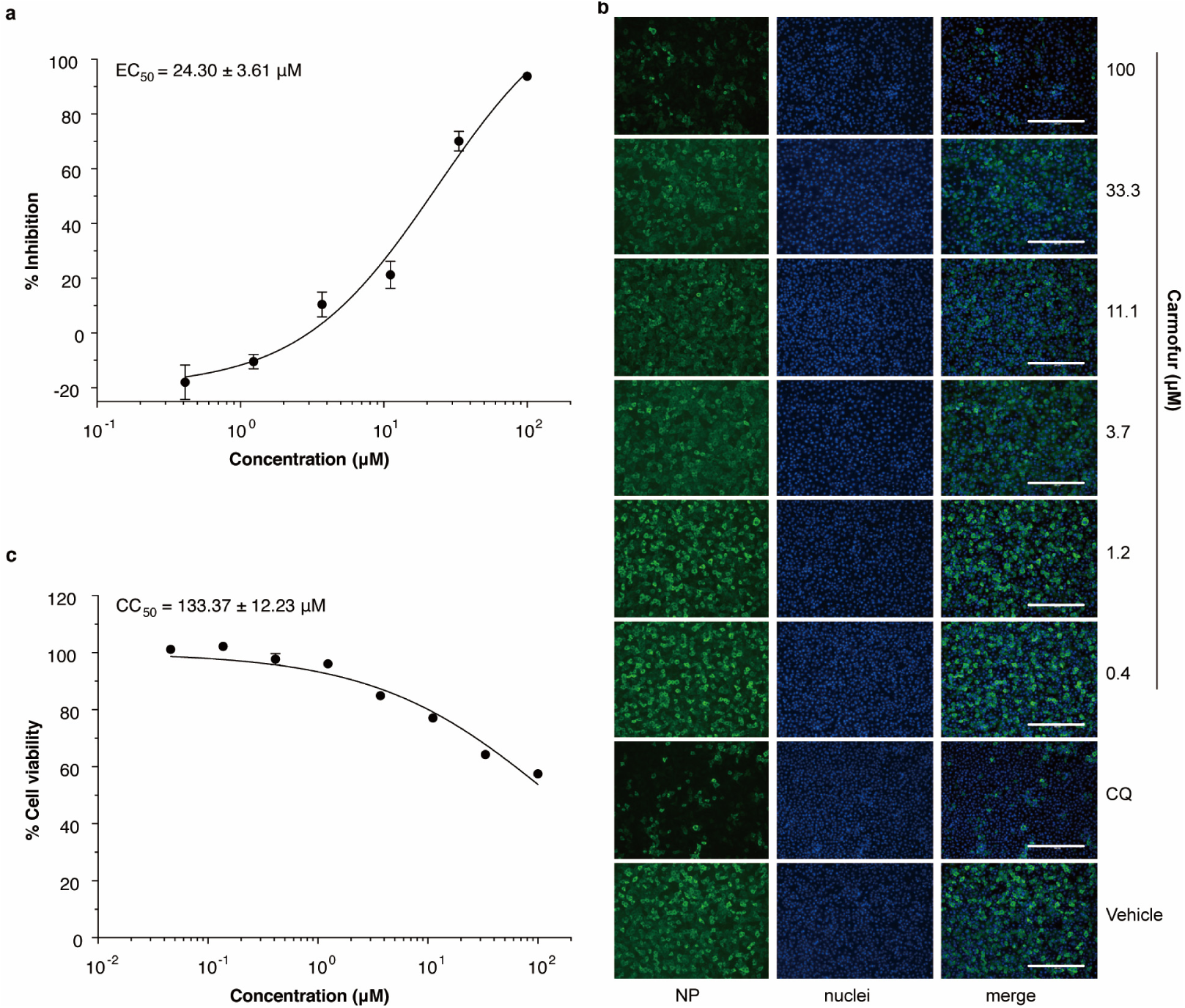
Inhibition of SARS-CoV-2 by Carmofur in Vero E6 cells. Vero E6 cells infected with SARS-CoV-2 at a MOI of 0.05 were treated with different concentrations of Carmofur. **a**, qRT-PCR assays were performed to measure viral copy number from cellular supernatant. The Y-axis of the graph indicates percentage inhibition of virus relative to DMSO (vehicle) treated sample. Data was shown as mean ± s.e.m., n = 6 biological replicates. **b**, Immunofluorescence for intracellular NP. At 24 hours post infection, cells were fixed, and intracellular NP levels were monitored by immunofluorescence. Chloroquine (CQ, 10 μM) was used as a positive control. Results are shown as representative of three biological replicates. Bars: 400 μm. **c**, Cell viability was measured using a Cell Counting Kit-8 (a commonly used kit for quantitation of viable cell number to measure proliferation and cytotoxicity) assay. The Y-axis represents percentage of cell viability relative to DMSO (vehicle) treated sample. Data was shown as mean ± s.e.m., n = 3 biological replicates. Data for graphs in **a** and **c** are available as source data.

In conclusion, the crystal structure of M^pro^ in complex with Carmofur shows that the compound directly modifies the catalytic Cys145 of SARS-CoV-2 M^pro^. Our study also provides a basis for rational design of Carmofur analogs with enhanced inhibitory efficacy to treat COVID-19. Since M^pro^ is highly conserved among all CoV M^pro^s, Carmofur and its analogs may be effective against a broader spectrum of coronaviruses.

## Methods

### Cloning, protein expression and purification of SARS-CoV-2 M^pro^

The cell cultures were grown and the protein expressed according to a previous report^7^. The cell pellets were resuspended in lysis buffer (20mM Tris-HCl pH 8.0, 150 mM NaCl, 5% Glycerol), lysed by high-pressure homogenization, and then centrifuged at 25,000g for 30 min. The supernatant was loaded onto Ni-NTA affinity column (Qiagen, Germany), and washed by the lysis buffer containing 20 mM imidazole. The His-tagged M^pro^ was eluted by lysis buffer that included 300 mM imidazole. The imidazole was then removed through desalting. Human rhinovirus 3C protease was added to remove the C-terminal His tag. SARS-CoV-2 M^pro^ was further purified by ion exchange chromatography. The purified M^pro^ was transferred to 10 mM Tris-HCl pH 8.0 through desalting and stored at -80 degrees until needed.

### Crystallization, data collection and structure determination

SARS-CoV-2 M^pro^ was concentrated to 5 mg/ml incubated with 0.3 mM Carmofur (Selleck, USA) for 1 hour and the complex was crystallized by hanging drop vapor diffusion method at 20 °C. The best crystals were grown using a well buffer containing 0.1 M MES pH 6.0, 5% polyethylene glycol (PEG) 6000, and 3% DMSO. The cryo-protectant solution was the reservoir but with 20% glycerol added.

X-ray data were collected on beamline BL17U1 at Shanghai Synchrotron Radiation Facility (SSRF) at 100 K and at a wavelength of 0.97918 Å using an Eiger X 16M image plate detector. Data integration and scaling were performed using the program XDS^21^. The structure was determined by molecular replacement (MR) with the PHASER^22^ and Phenix 1.17.1^23^ using the SARS-CoV-2 M^pro^ (PDB ID: 6LU7) as a search template. The model from MR was subsequently subjected to iterative cycles of manual model adjustment with Coot 0.8^24^ and refinement was completed with Phenix REFINE^25^. The inhibitor, Carmofur, was built according to the omit map. The phasing and refinement statistics are summarized in Supplementary Table 1.

### Antiviral and cytotoxicity assays for Carmofur

A clinical isolate of SARS-CoV-2 (nCoV-2019BetaCoV/Wuhan/WIV04/2019) was propagated in Vero E6 cells, and viral titer was determined as described previously^20^. Vero E6 cells were from ATCC with authentication. The authentication was performed by morphology check under microscopes and growth curve analysis. We confirm that all cells were tested as mycoplasma negative. For the antiviral assay, pre-seeded Vero E6 cells (5×10^4^ cells/well) were pre-treated with the different concentration of Carmofur for 1 h and the virus was subsequently added (MOI of 0.05) to allow infection for 1 h. Next, the virus-drug mixture was removed, and cells were further cultured with fresh drug containing medium. At 24 h post infection, the cell supernatant was collected and vRNA in supernatant was subjected to qRT-PCR analysis, while cells were fixed and subjected to immunofluorescence to monitor intracellular NP level as described previously^20^. For cytotoxicity assays, Vero E6 cells were suspended in growth medium in 96-well plates. The next day, appropriate concentrations of Carmofur were added to the medium. After 24 h, the relative numbers of surviving cells were measured by the CCK8 (Beyotime, China) assay in accordance with the manufacturer’s instructions. All experiments were performed in triplicate, and all the infection experiments were performed at biosafety level-3 (BSL-3).

### Reporting Summary

Further information on experimental design is available in the Nature Research Reporting Summary linked to this article.

### Data availability

Coordinates and structure factors for SARS-CoV-2 M^pro^ in complex with Carmofur have been deposited in Protein Data Bank (PDB) with accession number 7BUY. Source data for Fig. 2 are available with the paper online.

## Acknowledgements

We are grateful to the staff at the BL17U1, BL18U1 and BL19U1 at Shanghai Synchrotron Radiation Facility (SSRF, China), where data was collected. This work was supported by grants from National Key R&D Program of China (grants No. 2017YFC0840300 and 2020YFA0707500 to Z.R.), Project of International Cooperation and Exchanges NSFC (grant No. 81520108019 to Z.R.), Science and Technology Commission of Shanghai Municipality (grant No. 20431900200), Department of Science and Technology of Guangxi Zhuang Autonomous Region (grant No. 2020AB40007), and the Natural Science Foundation of China (grant No. 31970165).

## Author contributions

H.Y. and Z.R. conceived the project; Z.J., Y.Zhao, H.Y. and Z.R. designed the experiments; Z.J., Y.Zhao, H.W., Y.Zhu, C.Z., X.D., J.Y. and Xiuna Yang cloned, expressed, purified and crystallized proteins; Y.Zhao, Z.J., B.Z. and T.H. collected the diffraction data; Y.Zhao, B.Z. and X.L. solved the crystal structure; Y.S. and Y.W. performed cell-based antiviral and cytotoxicity assays; Y.D. and L.Z. performed qRT-PCR and cytotoxicity assay analysis; Z.J., Y.Zhao, Y.D., Xiaobao Yang, K.Y., X.L., L.W.G., G.X., L.Z., H.Y. and Z.R. analyzed and discussed the data; Z.J., Y.Zhao, K.Y., L.W.G., L.Z., H.Y. and Z.R. wrote the manuscript.

---

**Extended Data Fig. 1.**
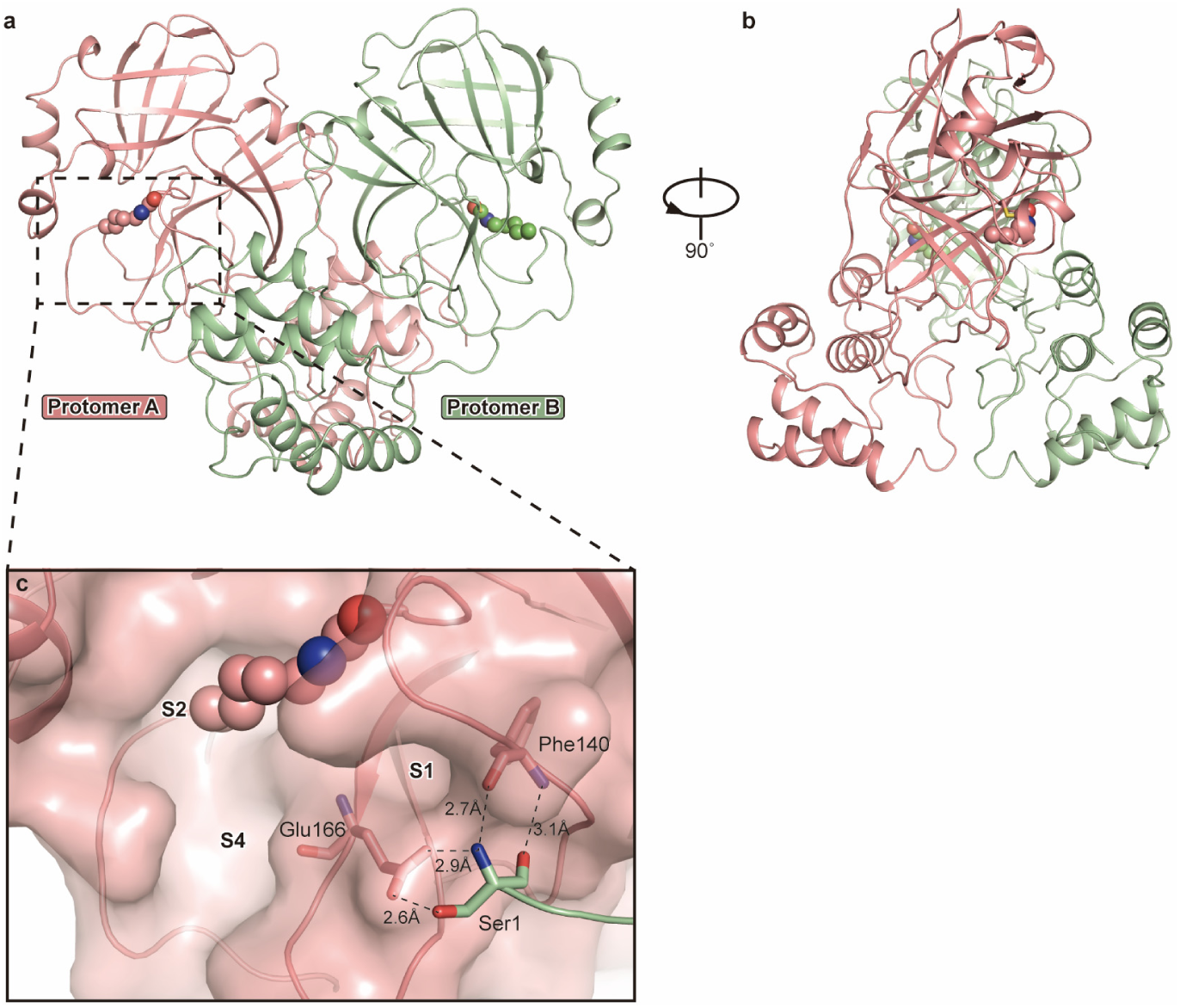
Overall structure of SARS-CoV-2 M^pro^ in complex with Carmofur. **a**, The overall structure of SARS-CoV-2 M^pro^ in complex with Carmofur. The salmon and green represent the different protomers. The Carmofur atoms are shown as solid spheres. **b**, The side view of the complex. **c**, The first serine participates in the formation of the dimer.

**Extended Data Fig. 2.**
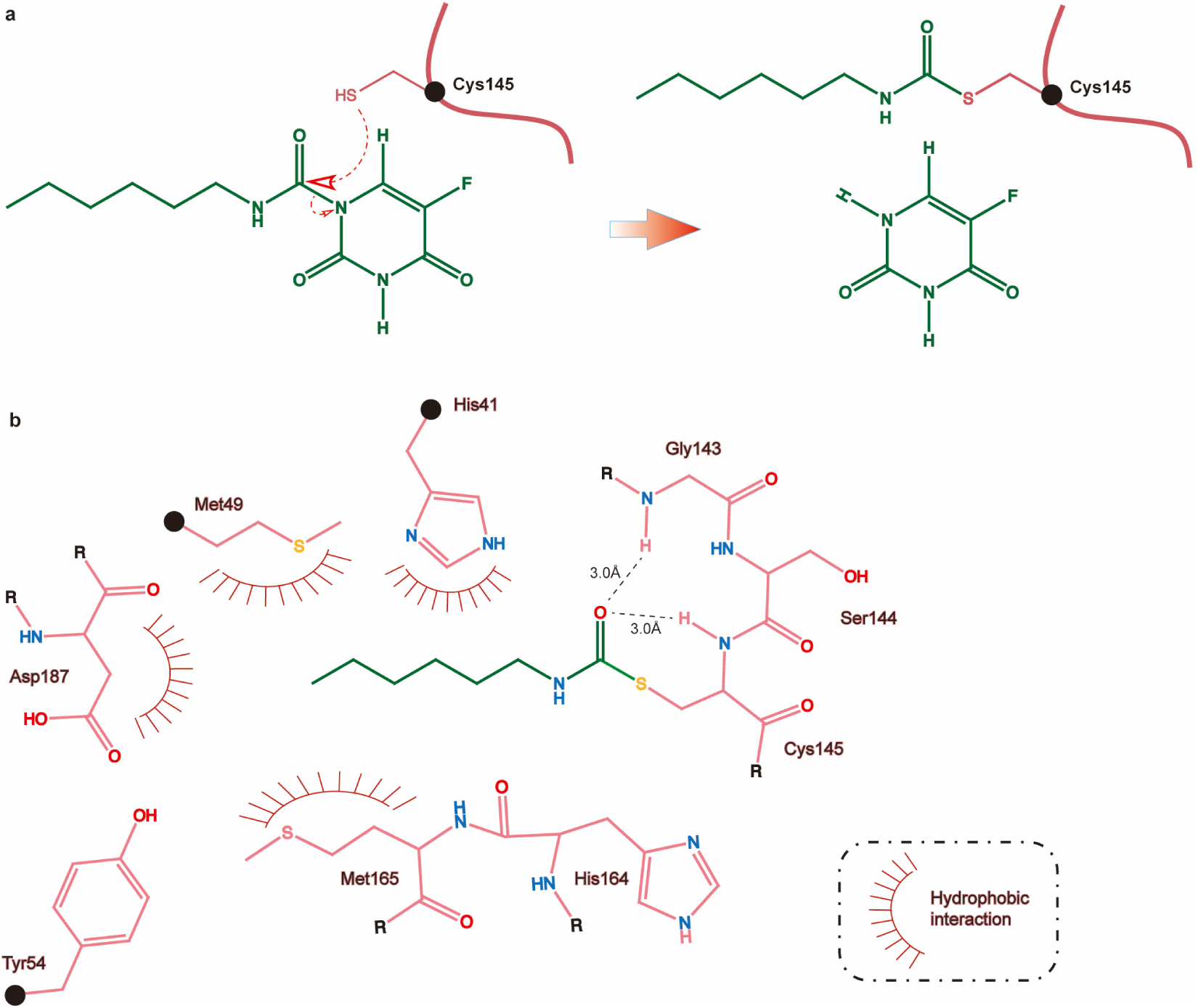
Inhibition of M^pro^ by Carmofur. **a**, Putative inhibition mechanism. Red curve represents M^pro^ polypeptide and the black sphere represent the C*α* of C145. **b**, Schematic diagram of M^pro^-Carmofur interactions. Black spheres represent C*α* atoms.

**Extended Data Fig. 3.**
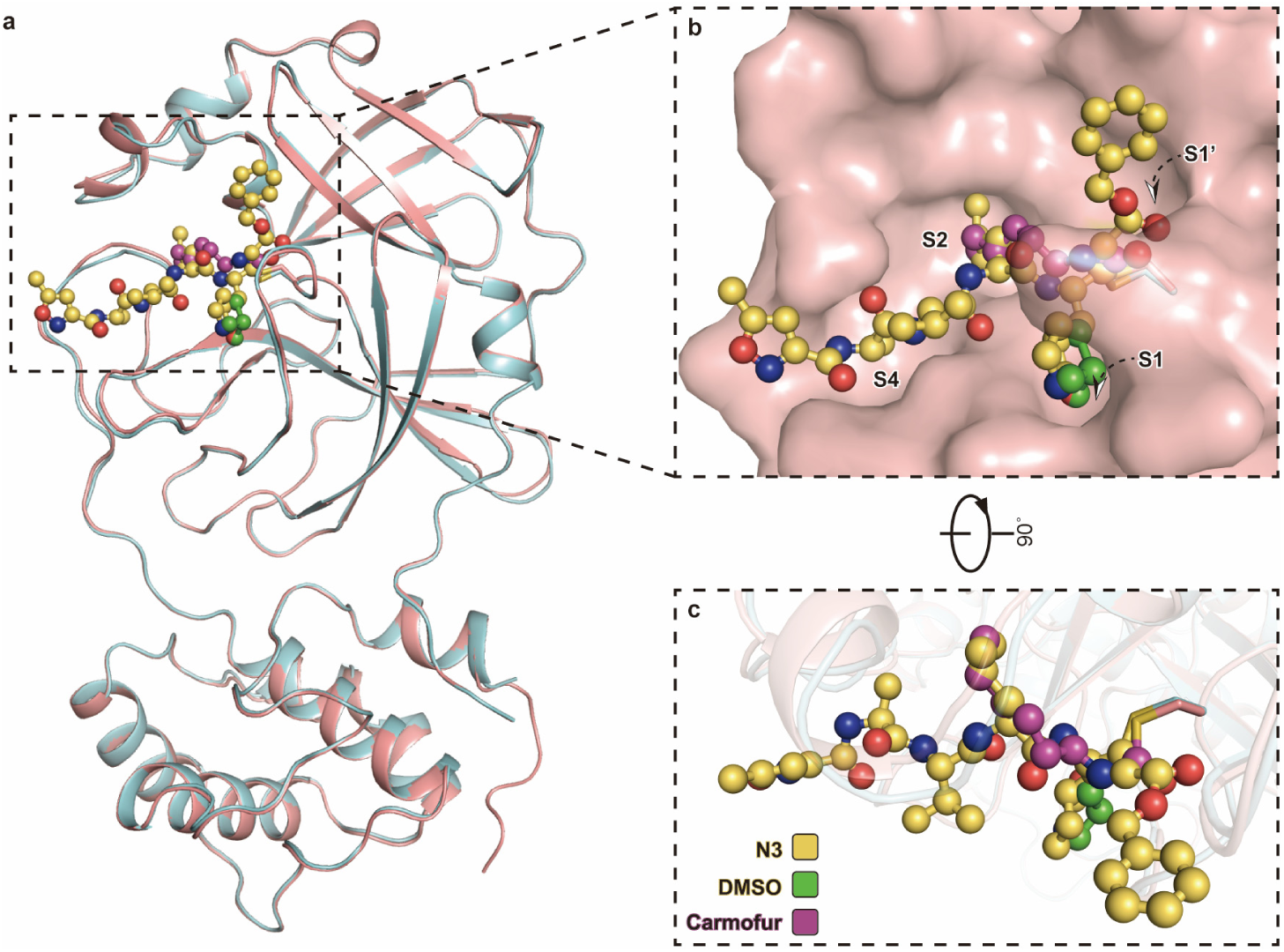
Binding mode of Carmofur and N3 to M^pro^. **a**, Overall structural comparison between the M^pro^-Carmofur and M^pro^-N3 complexes. The salmon cartoon represents the Carmofur bound structure and the light cyan represents the N3 bound structure. Carmofur, N3 and DMSO are represented by the purple, yellow and green balls and sticks, respectively. **b**, The binding pocket of M^pro^. Carmofur and N3 are represent in the same way as in panel **a. c**, Schematic diagram of Carmofur and N3.

**Supplementary Table 1.** Data collection and refinement statistics.

| PDB code: 7BUY |  |
| --- | --- |
| **Data Collection** |  |
| Space group | *C*2 |
| Cell dimensions |  |
| *a*, *b*, *c* (Å) | 98.03, 81.65, 51.64 |
| $\alpha$, $\beta$, $\gamma$ (°) | 90, 114.88, 90 |
| Resolution (Å) | 25.64 - 1.6 (1.66 - 1.60)<sup>a</sup> |
| *R*<sub>merge</sub> | 0.03824 (0.5052) |
| *R*<sub>meas</sub> | 0.04164 (0.5474) |
| *R*<sub>pim</sub> | 0.01622 (0.2089) |
| *I* / $\sigma I$ | 24.05 (3.60) |
| *CC*<sub>1/2</sub> | 0.999 (0.946) |
| Completeness (%) | 99.79 (99.96) |
| Redundancy | 3.4 (3.4) |
| **Refinement** |  |
| Resolution (Å) | 25.64-1.6 |
| No. reflections | 48563 (4843) |
| *R*<sub>work</sub> / *R*<sub>free</sub> | 0.1859 / 0.2034 |
| No. atoms |  |
| Protein | 2367 |
| Ligand/ion | 29 |
| Water | 205 |
| *B* factors |  |
| Protein | 37.91 |
| Ligand/ion | 52.55 |
| Water | 46.4 |
| R.m.s. deviations |  |
| Bond lengths (Å) | 0.016 |
| Bond angles (°) | 1.37 |
| Methods |  |
| Ramachandran (%) |  |
| Favored | 97.37 |
| Allowed | 2.63 |
| Outliers | 0 |

Data was collected from a single crystal. <sup>a</sup> Values in parentheses are for highest-resolution shell.

## References

1. Wu, F. et al. A new coronavirus associated with human respiratory disease in China. Nature 579, 265–269 (2020).

2. Zhou, P. et al. A pneumonia outbreak associated with a new coronavirus of probable bat origin. Nature 579, 270–273 (2020).

3. Hegyi, A. & Ziebuhr, J. Conservation of substrate specificities among coronavirus main proteases. J Gen Virol 83, 595–599 (2002).

4. Anand, K. et al. Structure of coronavirus main proteinase reveals combination of a chymotrypsin fold with an extra alpha-helical domain. Embo j 21, 3213–24 (2002).

5. Yang, H. et al. The crystal structures of severe acute respiratory syndrome virus main protease and its complex with an inhibitor. Proc Natl Acad Sci U S A 100, 13190–5 (2003).

6. Pillaiyar, T., Manickam, M., Namasivayam, V., Hayashi, Y. & Jung, S.H. An Overview of Severe Acute Respiratory Syndrome-Coronavirus (SARS-CoV) 3CL Protease Inhibitors: Peptidomimetics and Small Molecule Chemotherapy. J Med Chem 59, 6595–628 (2016).

7. Jin, Z. et al. Structure of Mpro from COVID-19 virus and discovery of its inhibitors. Nature (2020).

8. Sakamoto, J. et al. An individual patient data meta-analysis of adjuvant therapy with carmofur in patients with curatively resected colon cancer. Jpn J Clin Oncol 35, 536–44 (2005).

9. Morimoto, K. & Koh, M. Postoperative adjuvant use of carmofur for early breast cancer. Osaka City Med J 49, 77–83 (2003).

10. Gröhn, P. et al. Oral carmofur in advanced gastrointestinal cancer. Am J Clin Oncol 13, 477–9 (1990).

11. Nishio, S. et al. Study on effectiveness of carmofur (Mifurol) in urogenital carcinoma, especially bladder cancer, as a post-operative adjuvant chemotherapeutic agent. Hinyokika Kiyo 33, 295–303 (1987).

12. Ooi, A. et al. Plasma, intestine and tumor levels of 5-fluorouracil in mice bearing L1210 ascites tumor following oral administration of 5-fluorouracil, UFT (mixed compound of tegafur and uracil), carmofur and 5’-deoxy-5-fluorouridine. Biol Pharm Bull 24, 1329–31 (2001).

13. Sato, S., Ueyama, T., Fukui, H., Miyazaki, K. & Kuwano, M. Anti-tumor effects of carmofur on human 5-FU resistant cells. Gan To Kagaku Ryoho 26, 1613–6 (1999).

14. Nguyen, H.S., Awad, A.J., Shabani, S. & Doan, N. Molecular Targeting of Acid Ceramidase in Glioblastoma: A Review of Its Role, Potential Treatment, and Challenges. Pharmaceutics 10(2018).

15. Dementiev, A. et al. Molecular Mechanism of Inhibition of Acid Ceramidase by Carmofur. J Med Chem 62, 987–992 (2019).

16. Xue, X. et al. Structures of two coronavirus main proteases: implications for substrate binding and antiviral drug design. J Virol 82, 2515–27 (2008).

17. Ren, Z. et al. The newly emerged SARS-like coronavirus HCoV-EMC also has an “Achilles’ heel”: current effective inhibitor targeting a 3C-like protease. Protein Cell 4, 248–50 (2013).

18. Wang, F., Chen, C., Tan, W., Yang, K. & Yang, H. Structure of Main Protease from Human Coronavirus NL63: Insights for Wide Spectrum Anti-Coronavirus Drug Design. Sci Rep 6, 22677 (2016).

19. Yang, H. et al. Design of wide-spectrum inhibitors targeting coronavirus main proteases. PLoS Biol 3, e324 (2005).

20. Wang, M. et al. Remdesivir and chloroquine effectively inhibit the recently emerged novel coronavirus (2019-nCoV) in vitro. Cell Res 30, 269–271 (2020).

## References

21. Kabsch, W. XDS. Acta Crystallogr D Biol Crystallogr 66, 125–132 (2010).

22. McCoy, A.J. et al. Phaser crystallographic software. J Appl Crystallogr 40, 658–674 (2007).

23. Liebschner, D. et al. Macromolecular structure determination using X-rays, neutrons and electrons: recent developments in Phenix. Acta Crystallogr D Struct Biol 75, 861–877 (2019).

24. Emsley, P., Lohkamp, B., Scott, W.G. & Cowtan, K. Features and development of Coot. Acta Crystallogr D Biol Crystallogr 66, 486–501 (2010).

25. Afonine, P.V. et al. Towards automated crystallographic structure refinement with phenix.refine. Acta Crystallogr D Biol Crystallogr 68, 352–67 (2012).

